# Global signal regression reduces connectivity patterns related to physiological signals and does not alter EEG-derived connectivity

**DOI:** 10.1101/2024.04.18.590163

**Authors:** Alba Xifra-Porxas, Michalis Kassinopoulos, Prokopis Prokopiou, Marie-Hélène Boudrias, Georgios. D. Mitsis

**Author notes:** **Corresponding author:** Georgios D. Mitsis, Department of Bioengineering, McGill University. Montréal, Canada.

## Abstract

Functional brain connectivity measures extracted from resting-state functional magnetic resonance imaging (fMRI) scans have generated wide interest as potential noninvasive biomarkers. In this context, performing global signal regression (GSR) as a preprocessing step remains controversial. Specifically, while it has been shown that a considerable fraction of global signal variations is associated with physiological and motion sources, GSR may also result in removing neural activity. Here, we address this question by examining the fundamental sources of resting global signal fluctuations using simultaneous electroencephalography (EEG)-fMRI data combined with cardiac and breathing recordings. Our results suggest that systemic physiological fluctuations account for a significantly larger fraction of global signal variability compared to electrophysiological fluctuations. Furthermore, we show that GSR reduces artifactual connectivity due to heart rate and breathing fluctuations, but preserves connectivity patterns associated with electrophysiological activity within the alpha and beta frequency ranges. Overall, these results provide evidence that the neural component of resting-state fMRI-based connectivity is preserved after the global signal is regressed out.

## 1. Introduction

Studying the brain at rest has become a powerful tool towards revealing intrinsic characteristics of functional brain organization. Even in the absence of overt behaviour, brain activity fluctuates in an organized fashion in the form of large-scale brain networks that resemble those observed during behavioral tasks (Biswal et al., 1995; Fox and Raichle, 2007; Gratton et al., 2018; Smith et al., 2009). Functional magnetic resonance imaging (fMRI) has been widely used for studying the resting-state activity of temporally correlated and spatially distributed brain regions, and has shown promise to uncover mechanisms underlying neurological disorders (Gratton et al., 2019; Kassinopoulos et al., 2021; Xia et al., 2018). However, the blood-oxygenated level dependent (BOLD) signal typically employed in fMRI studies is only partially attributed to neural activity. Specifically, it has been shown that BOLD signal fluctuations are also driven by thermal noise as well as physiological and motion artifacts (Liu, 2016; Murphy et al., 2013).

The BOLD fMRI signal relies on changes in local cerebral blood flow (CBF) to detect neural activity, which constitutes an important caveat, as several additional processes may induce fluctuations in CBF. These include changes in heart rate and breathing patterns (Birn et al., 2006; Shmueli et al., 2007), variations in CO2 (Prokopiou et al., 2019; Wise et al., 2004), as well as arterial blood pressure (Whittaker et al., 2019a). Furthermore, the neurally driven fraction of the BOLD fMRI signal is not a direct measurement of neural activity, but an indirect measurement determined by neurovascular coupling mechanisms (Iadecola, 2017; Logothetis, 2008; Logothetis et al., 2001). It is also worth noting that the characteristics of physiological response functions that describe the dynamic effects of the aforementioned physiological signals (heart rate, respiration, arterial CO2 etc.) on the fMRI signal are highly similar to those of the hemodynamic response function (Prokopiou et al., 2022; Wu and Marinazzo, 2016), which further complicates disentangling the neural and physiological fMRI signal sources.

The aggregate nature of the BOLD fMRI signal described above renders the removal of its non-neural components a crucial and challenging task, which is particularly exacerbated in resting-state studies, where there is no *a priori* assumption for the temporal pattern of the underlying neural activity. Although there have been many advances in resting-state fMRI denoising (Caballero-Gaudes and Reynolds, 2017; Ciric et al., 2018), there is still no gold standard for resting-state fMRI preprocessing. In particular, global signal regression (GSR), which involves regressing out the average fMRI signal across the whole brain (global signal) from every voxel, has been proposed as a pre-processing step (Aguirre et al., 1998; Fox et al., 2009; Macey et al., 2004). Yet, as the processes underpinning the global signal are still poorly understood, GSR has turned out to be one of the most contentious preprocessing steps in fMRI denoising (Liu et al., 2017; Murphy and Fox, 2017; Power et al., 2017a). The rationale behind GSR is that the global signal mostly encompasses non-neuronal processes arising from physiological sources, head motion and scanner artifacts (Kassinopoulos and Mitsis, 2021; Power et al., 2017b). Therefore, GSR represents an effective data-driven approach for removing these global fluctuations, as they may result in artifactual functional connectivity patterns (Burgess et al., 2016; Ciric et al., 2017; Parkes et al., 2018; Xifra-Porxas et al., 2020). Indeed, GSR has been shown to increase the similarity of functional connectivity estimates across modalities (Keller et al., 2013). In addition, recent studies have reported that GSR improves the identifiability of well-established resting-state networks (Kassinopoulos and Mitsis, 2022), connectome fingerprinting accuracy (Xifra-Porxas et al., 2021), as well as the association between resting-state functional connectivity and behavioral measures (Li et al., 2019b), which suggests the potential benefits of performing GSR.

Yet, there is also converging evidence from simultaneous electrophysiological-fMRI studies that neural activity is linked to the global signal. Earlier investigations, albeit not directly examining the global signal, showed that fluctuations in local field potentials exhibited fairly widespread correlations with fMRI activity over the macaque brain (Scholvinck et al., 2010). More recently, fluctuations of the global signal have been linked to electrophysiological indices of arousal (C. W. Wong et al., 2016; Wong et al., 2013) and glucose metabolism (Thompson et al., 2016). Furthermore, global resting-state fluctuations were found to at least partially stem from the basal forebrain (Liu et al., 2018; Turchi et al., 2018). Gutierrez-Barragan and colleagues showed that brain states occur at specific phases of global fMRI signal fluctuations in the mouse brain (Gutierrez-Barragan et al., 2019). Finally, individual variations in global signal topography have been associated with behavioral measures (Li et al., 2019a). All these reports suggest that GSR may remove neuronal-related fluctuations of interest in functional connectivity studies.

Until now, the vast majority of studies investigating the processes underpinning the global signal have probed its physiological or neural origins separately. The neurally-related fraction of the global signal has been associated to fluctuations in arousal and vigilance, likely regulated by the autonomic nervous system (Oken et al., 2006; Olbrich et al., 2011). However, these apparent neural fluctuations could also be associated with changes in systemic physiological quantities such as heart rate and breathing, which can in turn influence the fMRI signal. Hence, there is potentially a closed loop path through which neural autonomic activity indirectly contributes to the global signal through changes in physiological signals. Overall, the above observations suggest that physiological and neural processes should be simultaneously taken into consideration to fully elucidate the nature of the resting-state global signal, as well as quantify the extent to which neural and physiological mechanisms generate similar or distinct contributions to global signal fluctuations.

While GSR is a more prevalent technique in resting-state studies, it was originally developed for task-based studies (Zarahn et al., 1997), and it is still frequently applied in this setting (e.g. (Finn et al., 2015; Marek et al., 2018)). Glasser et al. (2018) found that application of GSR to task fMRI data led to reduced statistical sensitivity for detecting activations, suggesting that the global signal in task-based studies contains relevant neural signal. However, the contributions of neurally-related fluctuations to the global signal during behavioral tasks have not yet been elucidated, as previous simultaneous electrophysiological-fMRI investigations of the global signal only examined resting-state conditions.

In the present study, we used simultaneously acquired electroencephalography (EEG)-fMRI data, as well as physiological recordings (heart rate and respiration), to quantify the unique and shared contributions of physiological and neural processes to the global fMRI signal, both at rest and during a hand-grip task. Furthermore, we generated two synthetic fMRI datasets that consisted of systemic and electrophysiological fluctuations, respectively, and evaluated the similarity between connectivity estimates extracted from the synthetic and experimental fMRI datasets. This allowed us to examine the effects of GSR on connectivity estimates and address explicitly whether eliminating the bias introduced by physiological processes inadvertently removes connectivity patterns related to electrophysiological activity.

## 2. Materials and methods

### 2.1 Participants

A total of 12 healthy volunteers (25.1 ± 2.9 years; 4 female) participated in the study. All subjects were right-handed according to the Edinburgh Handedness Inventory (Oldfield, 1971) and had no history of neurological or psychiatric disorders. The study was approved by the McGill University Ethical Advisory Committee. All participants signed a written informed consent and were compensated for their participation.

### 2.2 Experimental paradigm

The protocol carried out inside the MR scanner consisted of two 15-min resting-state runs, with a 13-min hand-grip fMRI task interleaved between them (Figure 1a). During the resting-state periods, subjects were instructed to stare at a white fixation cross displayed on a dark background and not to think of anything in particular. After the first rest period, the maximum voluntary contraction (MVC) was obtained for each participant, using the same hand gripper subsequently employed for the motor task. The motor task was a unimanual isometric right hand-grip, during which the subjects had to apply force to track a ramp target as accurately as possible (Larivière et al., 2019; Xifra-Porxas et al., 2019). At the onset of the trial, an orange circle appeared on the screen, and the subjects had 2 s to increase their force to reach a white target block at 15% of their MVC. This force was held for 3 s. Following this, participants tracked a linear increase of the force to reach 30% of their MVC over a 3-s period, during which they had to maintain the circle inside the white target block, followed by a 3-s hold at 30% of their MVC (Figure 1b). A single trial lasted 11 s and the inter-trial interval was jittered between 3 and 5 s, during which subjects stared at a white cross. The task consisted of 50 trials, resulting in a total duration of about 13 min.

**Figure 1.**
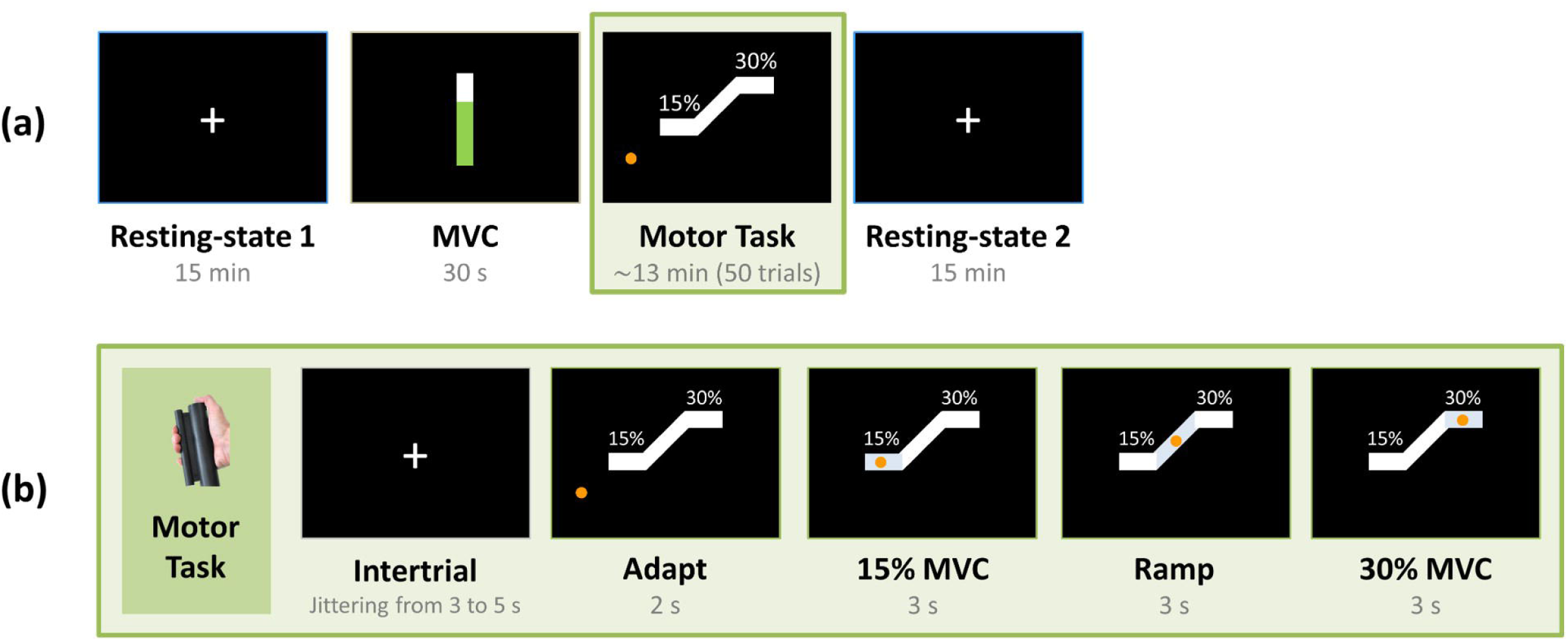
(a) Experimental paradigm. Participants underwent two resting-state scans with eyes open, alternated by a motor task. The maximum voluntary contraction (MVC) of each participant was measured before performing the motor task. **(b) Hand-grip task.** Participants performed a unimanual right hand-grip task. During each trial, they initially fixated on a crosshair for a few seconds. This was followed by the appearance of an orange circle on the screen, whereby participants had 2 s to apply force to reach 15% of their MVC. A steady grip was then maintained for 3 s, which was followed by a guided ramp period where participants had to apply force to reach 30% of their MVC and sustain this grip strength for another 3 s.

### 2.3 Data acquisition: EEG-fMRI data and physiological recordings

All experiments were conducted at the McConnell Brain Imaging Centre (BIC) of the Montreal Neurological Institute (MNI), McGill University, using a 3T Siemens MAGNETOM Prisma fit MRI scanner (Siemens AG, Germany). A 32-channel head coil was used to acquire whole-brain T2*-weighted functional gradient echo planar (EPI) data (3×3×4 mm^3^ voxels, TR=2120 ms, TE=30 ms, flip angle=90°, FOV=192×192 mm^2^, anterior-posterior phase encoding direction). Volumes were recorded in 35 transverse slices in descending order. A high-resolution structural volume was also acquired using a T1-weighted magnetization-prepared rapid acquisition gradient echo (MPRAGE) sequence (TR=2300 ms, TE=2.32 ms, flip angle=8°, FOV=240×240 mm^2^, 0.9 mm^3^ isotropic voxels).

EEG data were simultaneously recorded with an MR-compatible 64-channel system with Ag/AgCl ring-type electrodes (BrainAmp MR, Brain Products GmbH, Germany), sampled at 5000 Hz. Electrode impedances were maintained below 20 kΩ and verified both during setup and at the end of each scanning session. An equidistant electrode layout was used with AFz and Cz as ground and online recording reference, respectively. The EEG acquisition clock was synchronised with the MR scanner clock through a device that sent triggers to the EEG recording system every time an fMRI volume was acquired (TriggerBox, Brain Products GmbH, Germany). The electrodes were precisely localized using a 3-D electromagnetic digitizer (Polhemus Isotrack, USA).

Cardiac and breathing measurements were continuously recorded throughout the experiment using a pulse oximeter and respiratory belt (BIOPAC Systems, Inc., USA). An MR-compatible hand clench dynamometer (BIOPAC Systems, Inc., USA) was used to measure the subjects’ hand-grip strength during the motor paradigm. The pulse oximeter, respiratory belt and dynamometer were connected to an MP150 data acquisition system (BIOPAC Systems, Inc., USA) from which the signals were transferred to a computer, sampled at 1000 Hz.

### 2.4 Preprocessing

#### 2.4.1 fMRI data

The fMRI data were preprocessed using FSL (Jenkinson et al., 2012). Automated brain extraction using BET, motion correction via volume realignment using MCFLIRT, spatial smoothing (5 mm FWHM Gaussian kernel) and high-pass temporal filtering (100 s cutoff) were performed. Additional preprocessing included motion censoring based on the frame-wise displacement (FD) and root mean square intensity change of BOLD signal across the whole brain (DVARS) measures (Power et al., 2012). Motion censoring was applied by discarding volumes with FD>0.25 mm or when DVARS exceeded its median absolute deviation by a factor of 3, as well as their adjacent volumes. Volumes with subthreshold values of FD and DVARS were also discarded if they were preceded and followed by flagged volumes. All scans from one participant were excluded from the analysis, since more than 40% of volumes were identified as being contaminated by motion (Suppl. Fig. 1). Therefore, results from a total of 11 subjects are presented below.

The censored fMRI data were co-registered to each subject’s structural image and normalized to MNI space (2 mm). Subsequently, the preprocessed data were parcellated into 300 regions of interest (ROIs) using the Seitzman atlas (Seitzman et al., 2020). In that work, ROIs were assigned to large-scale functional networks using the InfoMap community detection algorithm (Rosvall and Bergstrom, 2008), applied to resting-state functional connectivity data. The fifteen ROIs that were not assigned to any large-scale network in Seitzman et al. (2020) were excluded here from further analysis, resulting in a final set of 285 ROIs.

We examined three sets of experimental fMRI data that differed in the set of nuisance regressors removed:

i. minimally preprocessed (MinPrep) fMRI data, with no additional denoising;
ii. preprocessed fMRI data where white matter denoising was applied; and
iii. preprocessed fMRI data with global signal regression (GSR), where both white matter denoising and GSR was performed.

In the second dataset, white matter denoising was performed using principal component analysis (PCA) on the parcellated data to mitigate the effects of head motion not corrected by motion censoring as well as the effects of cardiac pulsatility and breathing-related motion (Behzadi et al., 2007; Kassinopoulos and Mitsis, 2022; Muschelli et al., 2014). Specifically, 10 white matter PCA components were removed from the data, as this number was found to maximize the functional connectivity contrast (FCC; Figure 2), which is a quality control metric for evaluating the identifiability of well-established large-scale networks (Kassinopoulos and Mitsis, 2022). In the third dataset, the global signal and the same 10 white matter PCA components were simultaneously regressed out.

**Figure 2.**
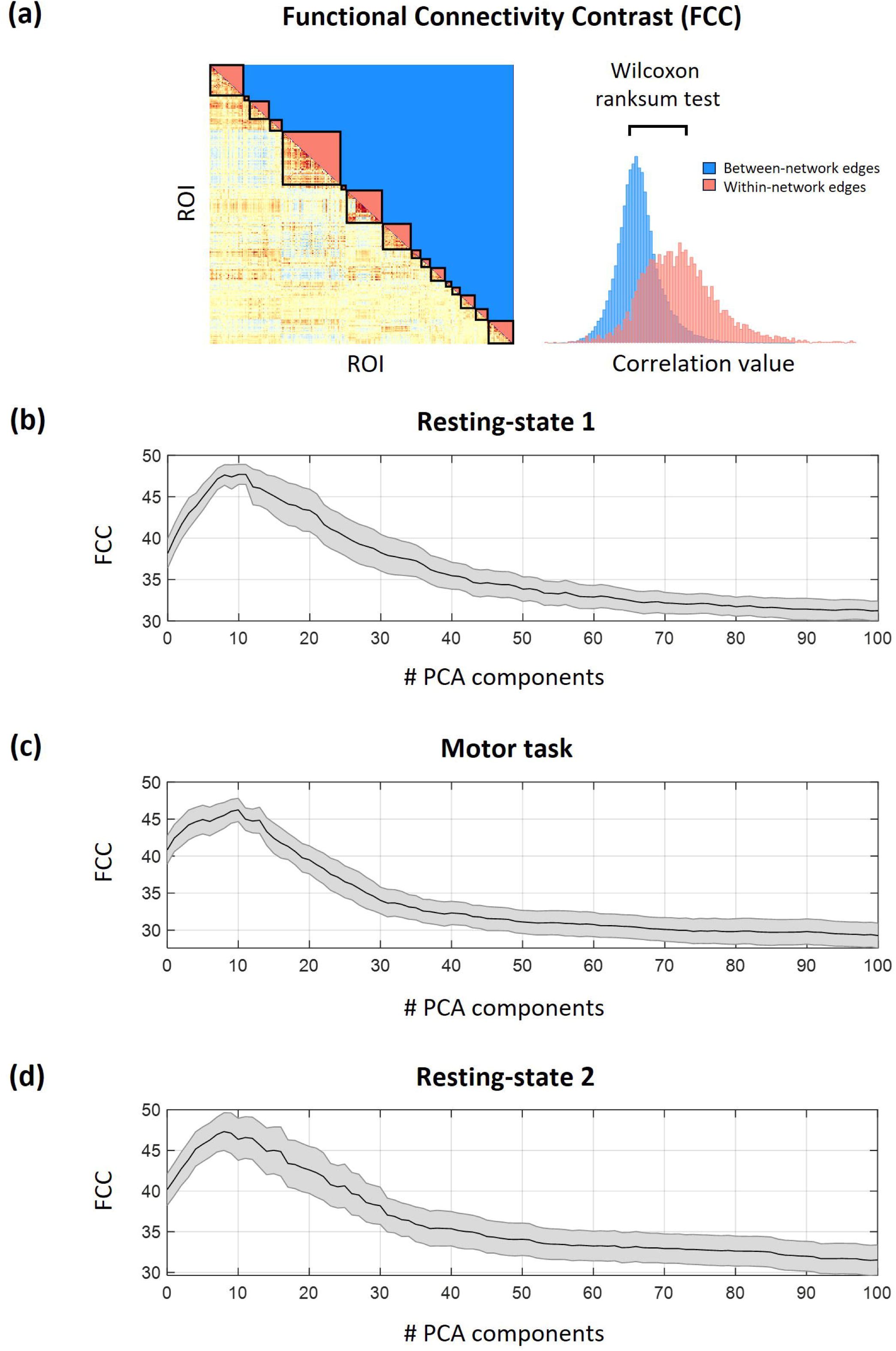
Optimal number of PCA white matter components used for fMRI denoising. **(a)** Calculation of the functional connectivity contrast (FCC) measure, which quantifies the difference in correlation values among edges within and between-networks and serves in assessing the identifiability of well-established large-scale networks. **(b-d)** Group-averaged FCC measure as a function of the number of white matter PCA components used for denoising for **(b)** resting-state 1, **(c)** motor task, and **(d)** resting-state 2. The black line denotes mean across subjects and the shaded area denotes standard error. Based on these traces, 10 PCA components were used for denoising, as this number yielded a high FCC value for all three conditions.

#### 2.4.2 EEG data

EEG data were corrected offline for gradient and ballistocardiogram (BCG) artifacts using sliding average template subtraction in BrainVision Analyzer 2 (Brain Products GmbH, Germany). The data were subsequently downsampled to 200 Hz and further preprocessed using independent component analysis to remove remaining non-neural components such as ocular and muscle artifacts, as well as gradient and BCG residuals. Non-neural components were identified through visual inspection by three of the paper’s authors (AXP, MK, PP), based on classification guidelines provided in the ICLabel tutorial (https://labeling.ucsd.edu/tutorial/labels). The preprocessed data were re-referenced using the average signal from all channels.

Time-frequency spectrograms were calculated for each EEG channel using the continuous wavelet transform (CWT) with analytic Morlet wavelets in Matlab environment. Matlab’s default settings were used, which apply logarithmically spaced frequencies and 10 voices per octave. The frequency range was automatically determined based on the signal duration and sampling rate (i.e. 200 Hz). A global spectrogram was then obtained by calculating the root mean square across all channel-specific spectrograms. Then, we extracted the instantaneous power time-series through averaging of the global spectrogram within the following frequency bands: delta (1.5-4 Hz), theta (4-8 Hz), alpha (8-15 Hz) and beta (15-26 Hz). These global EEG power time-series were subsequently convolved with a canonical double-gamma haemodynamic response function and downsampled to the frequency of image volume sampling (TR=2.12 s). The censored frames flagged during fMRI preprocessing were removed from the convolved EEG power time-series.

#### 2.4.3 Physiological recordings

Beat-to-beat intervals were detected from the pulse oximeter signal, and the heart rate signal was computed as the inverse of the time differences between pairs of adjacent peaks and converted to beats-per-minute (bpm). Heart rate traces were visually examined to identify outliers, and an outlier replacement filter was used to eliminate spurious changes in heart rate. The breathing signal from the respiratory belt was detrended linearly, visually inspected and corrected for outliers using a replacement filter, low-pass filtered at 5 Hz, and z-scored. The respiratory flow, proposed in Kassinopoulos and Mitsis (2019) as a robust measure of the absolute flow of inhalation and exhalation of a subject at each time point, was extracted by further smoothing the breathing signal (moving average filter using a 1.5 sec window) and computing the square of the derivative of the smoothed breathing signal. Both heart rate and respiratory flow signals were resampled to 10 Hz.

Following this, we modelled the effect of systemic low-frequency oscillations (SLFOs) associated with heart rate and breathing patterns using a recently developed method that computes scan-specific physiological response functions (PRFs) (Kassinopoulos and Mitsis, 2019; scripts available on https://github.com/mkassinopoulos/PRF_estimation/). Briefly, this algorithm estimates PRF curves so that the convolution between heart rate and respiratory flow with their corresponding PRFs optimizes the fit on the global signal of the same scanning session, while ensuring that the shapes of the PRF curves are physiologically plausible. The heart rate and respiratory flow were convolved with their respective PRFs and added to obtain a time-series that reflects the total effect of SLFOs.

It is important to note that SLFOs reflect slow physiological modulations occurring at low frequencies up to ∼0.15 Hz, such as fluctuations in heart rate and breathing patterns. These processes unfold over time scales of ∼10 seconds and are well captured by our sampling rate [1/TR = 1/(2.120 s) ≈ 0.47 Hz], which satisfies the Nyquist criterion. While cardiac pulsatility artifacts occur at higher frequencies (∼1 Hz) and are subject to aliasing at typical TRs, our study does not aim to model such high-frequency effects. Instead, the PRF-based approach targets the slower systemic fluctuations that influence the BOLD signal more gradually and are not significantly impacted by the TR used in our acquisition.

### 2.5 Data analysis

We initially quantified the variance of the fMRI global signal explained by SLFOs and EEG power using partial correlation. Specifically, the contribution of SLFOs to the global signal was computed controlling for the EEG power time-series and vice versa. To assess the significance of these contributions to the global signal, surrogate data were generated via inter-subject surrogates (Lancaster et al., 2018), using physiological and EEG power time-series from different subjects. The contributions obtained using the experimental data from the same subject were compared to the ones obtained from the surrogate data using the Wilcoxon rank-sum test. The significance level was set to 0.05, and the p-values for the EEG bands were adjusted for multiple comparisons using the false discovery rate (FDR) method.

In addition, we generated three synthetic fMRI datasets, namely the SLFOs synthetic fMRI, alpha power synthetic fMRI, and beta power synthetic fMRI, following the framework employed in Xifra-Porxas et al. (2021). In brief, the ROI fMRI time-series *y*(*t*) in each of the 285 atlas-based ROIs was modelled using multiple linear regression, as follows:

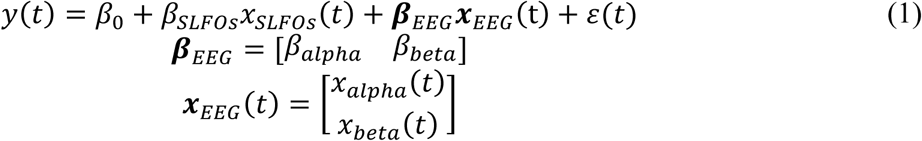

where *x*_*SLFOs*_(*t*) is the physiological regressor related to SLFOs, ***x***_*EEG*_(*t*) are the two EEG power time-series, {*β*_0_, *β*_*SLFOs*_, ***β***_*EEG*_} are the unknown parameters estimated for each of the three experimental datasets, and *ε* is the error.

Then, for each of the three processes of interest (i.e. SLFOs, alpha and beta EEG power), the estimated value *β̂* was multiplied by its corresponding regressor to obtain the associated fluctuations (*ŷ*_*PI*_ (*t*)) within a specific ROI, and a ‘clean’ time-series (*ŷ*_*PI*+𝑟𝑒𝑠_(*t*)) was calculated via removal of all other regressors, as follows:

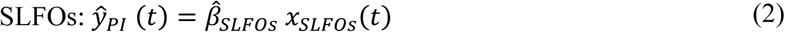

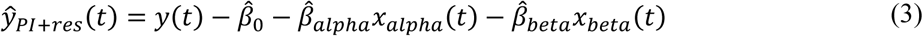

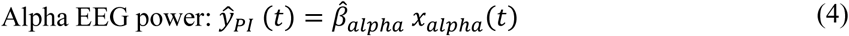

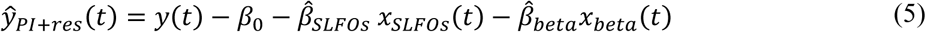

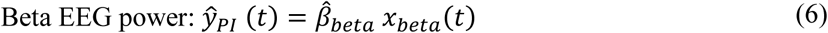

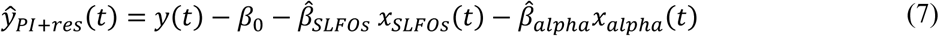

In this manner, we generated ‘cleaned’ ROI time-series (*ŷ*_*PI*+𝑟𝑒𝑠_(*t*)) in which all regressors were removed except the one corresponging to the specific process being evaluated. Subsequentely, the contribution of the latter to the remaining fluctuations within each ROI was quantified. To achieve this, the estimated process signal was correlated to the ‘clean’ ROI time-series:

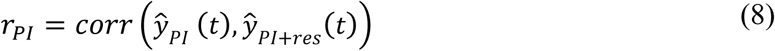

Afterwards, the estimated process signal was subtracted from the ‘clean’ ROI time-series to obtain the ‘residual’ time-series. These time-series were correlated to the ‘clean’ time-series to quantify the contribution of the ‘residual’ variations to the ‘clean’ ROI time-series:

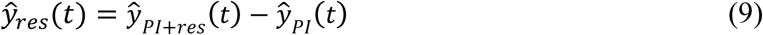

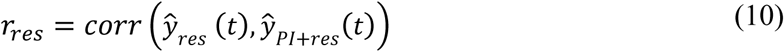

Finally, synthetic datasets were constructed for each process by combining the normalized process-specific signal and a first-order autoregressive (AR(1)) process (𝜓(*t*)) scaled by their respective correlation coefficients:

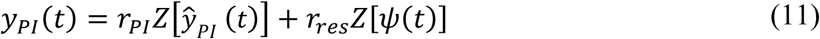

where Z[·] denotes normalization to zero mean and unit variance.

Using the framework described above, we constructed synthetic fMRI datasets in which the fluctuations attributed to the process of interest within each ROI was preserved while the remaining fraction of fluctuations was replaced with uncorrelated random signal of equal amplitude (Xifra-Porxas et al., 2021). We only considered the alpha and beta power among the EEG bands because they were found to be the only ones contributing to the global signal (Figure 3; *p* < 0.05).

**Figure 3.**
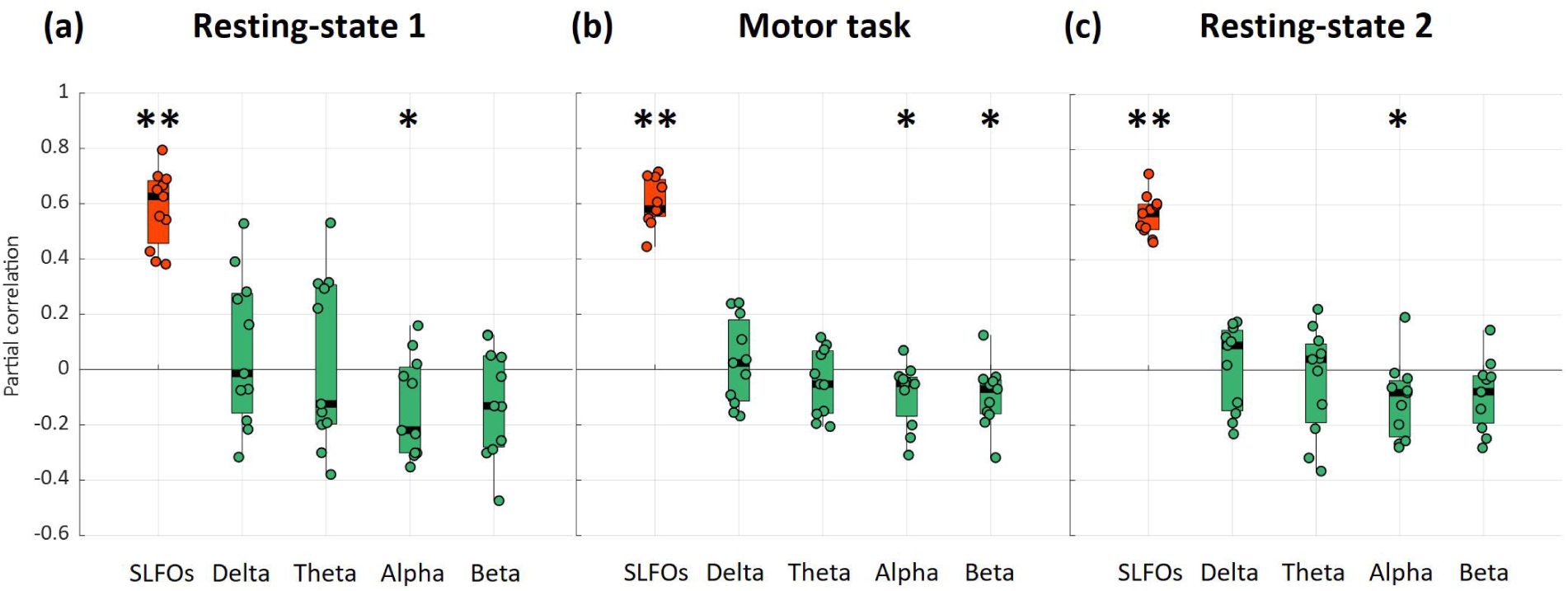
Partial correlation of SLFOs and EEG power time-series with the fMRI global signal across subjects for (a) resting-state 1, (b) motor task, and (c) resting-state 2. Significance testing was performed against surrogate data (Wilcoxon rank-sum test, * *p* < 0.05, ** *p* < 0.001). The p-values for the four EEG bands were corrected for multiple comparisons using FDR. SLFOs were highly correlated with the global signal, whereas alpha and beta power fluctuations were found to be weakly correlated with global signal variations.

We then computed the static functional connectivity matrices (i.e. pairwise correlations between time-series of all ROIs) for each of the three experimental datasets described earlier: minimally preprocessed experimental fMRI, preprocessed experimental fMRI with white matter denoising, and preprocessed experimental fMRI with both white matter denoising and GSR; as well as for the three synthetic datasets: SLFOs synthetic fMRI, alpha power synthetic fMRI, and beta power synthetic fMRI. Finally, we quantified the similarity between the SLFOs, alpha power and beta power connectivity matrices with respect to the fMRI connectivity matrices derived from the three experimental datasets, by computing the correlations between their upper triangular values. High similarity is expected when a process, whether SLFOs or EEG band power, explains a substantial portion of BOLD variance across multiple ROIs, resulting in elevated inter-regional correlations among those regions and imposing a spatial covariance structure reflected in both the synthetic and empirical functional connectivity matrices. The derived correlations were utilized to assess the effect of GSR on the similarity between SLFOs and fMRI connectivity, as well as EEG power and fMRI connectivity.

## 3. Results

### 3.1 Association of SLFOs and EEG power with the fMRI global signal

All signals examined (i.e. fMRI global signal, SLFOs, and EEG band power signals) exhibited prominent low-frequency fluctuations below 0.15 Hz, as shown in Supplementary Figure 2. In line with previous studies (Birn et al., 2006; Kassinopoulos and Mitsis, 2019; Power et al., 2017a; Shmueli et al., 2007), SLFOs were strongly correlated with the global signal, both at rest and during the motor task (Figure 3). Regarding the EEG bands, alpha power was negatively correlated with the global signal both at rest and during the motor task (Figure 3), and beta power was negatively correlated with the global signal during the motor task (Figure 3b). Illustrative individual time series from representative subjects can be found in Suppl. Fig. 2. It is worth noting that the correlation strength of alpha and beta fluctuations with the global signal, albeit significant, was considerably weaker compared to the correlation between SLFOs and the global signal. The mean correlation value across subjects for the full model containing both SLFOs and significant EEG power bands was equal to 0.64 ± 0.12 (resting-state 1), 0.64 ± 0.09 (motor task), and 0.59 ± 0.09 (resting-state 2). Moreover, alpha power fluctuations were found to be negatively correlated with SLFOs (𝑅 = −0.09, 𝑝 = 0.02) during resting-state 1. This correlation vanished when alpha fluctuations and SLFOs were controlled for global signal fluctuations (R=0.02, *p*=0.84), which may suggest a common component between those two variables that is reflected on the global signal, possibly reflecting a direct or indirect effect of autonomic activity. Yet, we were unable to replicate this result during the motor task or resting-state 2 (i.e. alpha power was not significantly correlated with SLFOs). Finally, delta and theta power fluctuations did not yield a consistent contribution to the global signal across subjects for any of the scans.

### 3.2 Effect of GSR on connectome patterns associated with SLFOs and EEG power fluctuations

For each subject and scanning session, connectivity matrices were derived using the synthetic dataset generated from SLFOs, enabling us to identify the connectivity edges more prone to exhibit artifactual connectivity attributable to SLFOs. We found that fluctuations due to SLFOs artifactually increased the connectivity across most brain networks and particularly within the visual network, as well as between the visual network and other brain networks (Figure 4a), consistent with previous observations using a larger cohort (Xifra-Porxas et al., 2021). Furthermore, the effect of SLFOs on the visual network were found to be spatially homogeneous at rest, whereas during the motor task, some edges were more affected than others (Figure 4a). Notably, the contribution of SLFOs to functional connectivity appeared to be attenuated during the second resting-state session compared to the first, suggesting session-related variability in SLFOs-induced artifacts. Consistent with our earlier study (Xifra-Porxas et al., 2021), WM denoising led to a significant reduction in similarity between the SLFOs’ and the fMRI connectivity matrices both at rest and during the motor task (Figure 4b; *p* < 0.001). Further, GSR yielded a small additional reduction in similarity for all three scans (*p* < 0.05). This result supports the extensive evidence that, at least with respect to mitigating nuisance processes, GSR is effective.

**Figure 4.**
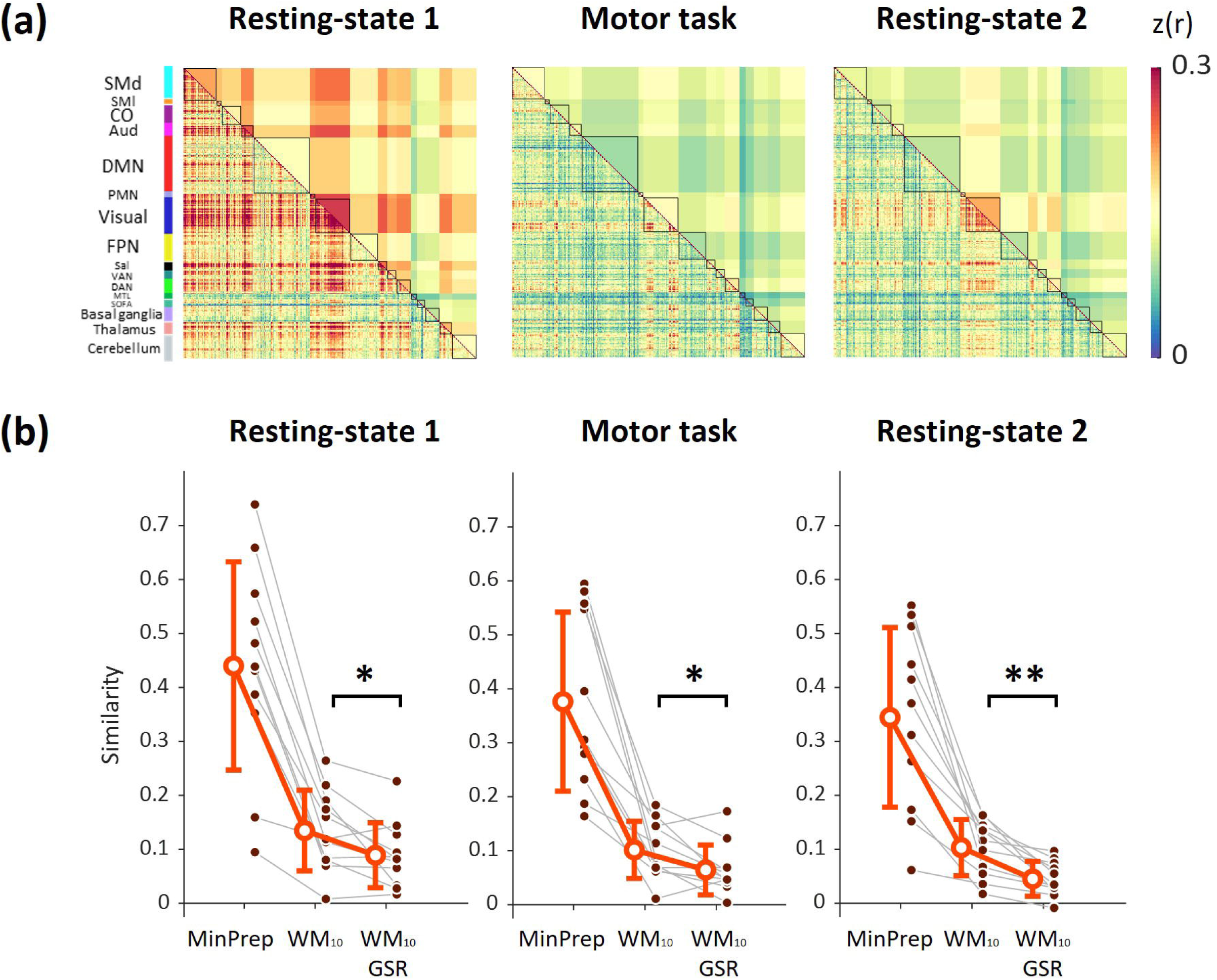
Effect of GSR on connectome patterns induced by SLFOs. **(a)** Group averaged FC matrices computed using the synthetic dataset associated with systemic low frequency fluctuations (SLFOs) for each scanning session. **(b)** Similarity between the SLFOs’ FC matrices and the FC matrices extracted from the minimally preprocessed (MinPrep) data, the preprocessed data (after regressing out 10 PCA white matter components; WM10), and the preprocessed data after global signal regression (WM10 & GSR). While WM regression reduced the similarity between the minimally preprocessed (MinPrep) and SLFOs connectivity matrices, suggesting a reduction in connectivity bias, GSR further reduced this similarity significantly (Wilcoxon rank-sum test, * *p* < 0.05, ** *p* < 0.005). Error bars denote standard deviation.

Subsequently, we sought to determine whether applying GSR also removed substantial EEG power fluctuations. First, we derived connectivity matrices using the fMRI synthetic datasets generated using the BOLD fluctuations attributed to alpha and beta power activity, thus highlighting the edges underpinning connectivity patterns of likely neural origin. At rest, alpha power variations mostly contributed to connectivity within the visual network and dorsal attention network as well as between these two networks (Figure 5a). On the other hand, during the motor task, alpha power variations mostly contributed to connectivity within the default mode network (Figure 5a), whereas beta power variations mainly contributed to connectivity within some edges of the visual network (Figure 5c).

**Figure 5.**
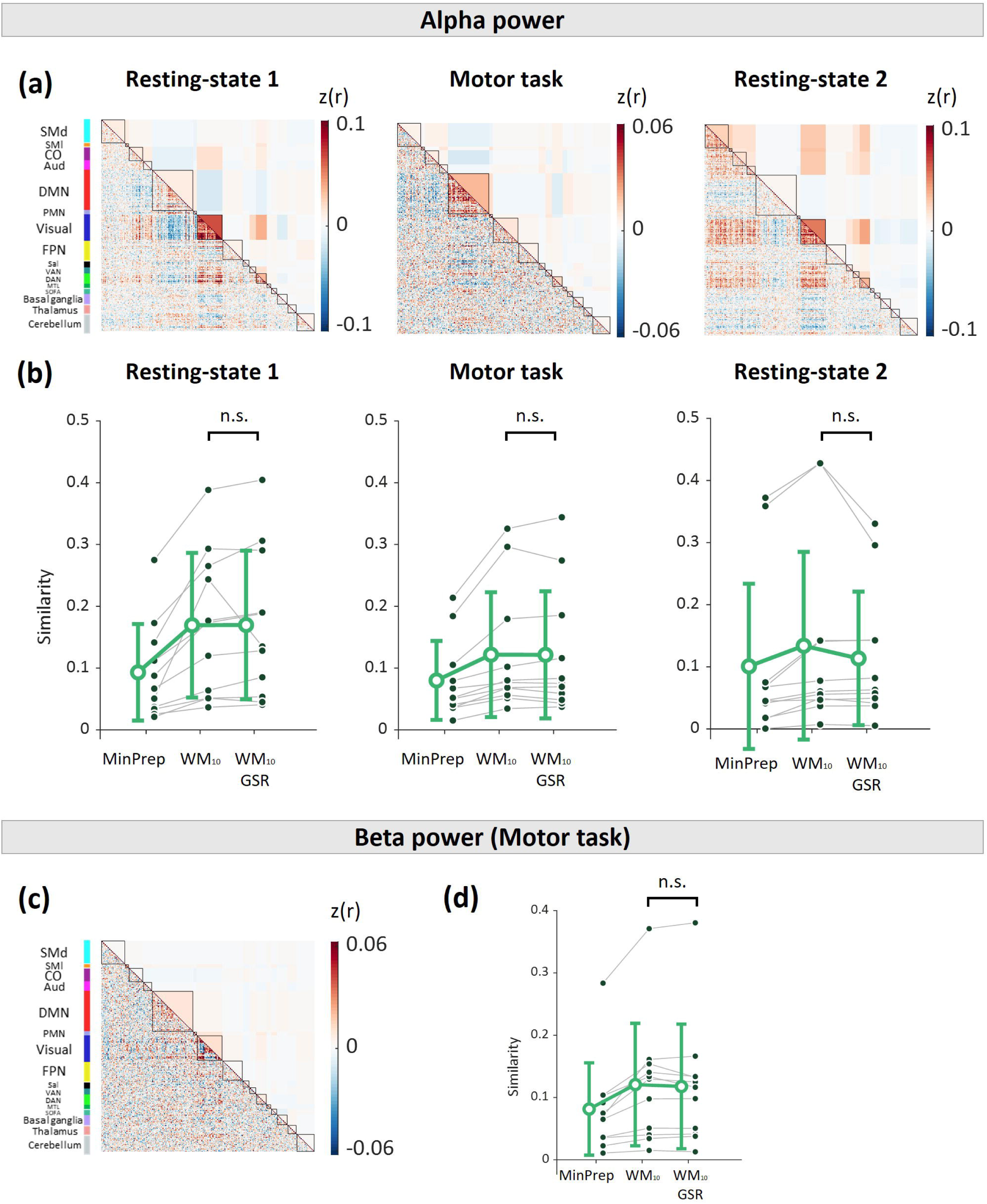
Effect of GSR on connectome patterns associated with EEG power. **(a)** Group averaged FC matrices computed using the fMRI synthetic dataset associated with fluctuations in alpha power, for each scanning session. **(b)** Similarity between the alpha band EEG-based FC matrices and the FC matrices extracted from the minimally preprocessed data, the preprocessed data (i.e. fMRI data after regressing out 10 PCA white matter components; WM10), and the preprocessed data after global signal regression (WM10 & GSR). **(c)** Group averaged FC matrices computed using the fMRI synthetic dataset associated with fluctuations in beta power, for the motor task. **(d)** Similarity between the beta band EEG-based FC matrices and the FC matrices extracted from the minimally preprocessed data (MinPrep), the preprocessed data (WM10), and the preprocessed data after GSR (WM10 & GSR). GSR did not significantly alter the connectivity related to EEG activity (Wilcoxon rank-sum test). In all subplots, error bars denote standard deviation.

Finally, we evaluated whether performing GSR on the preprocessed fMRI data significantly removed the connectivity signature of these neural processes. To do this, we compared the similarity of the connectome patterns associated with EEG activity with the connectivity patterns of the three experimental fMRI datasets. WM denoising was found to increase the similarity between modalities for all scans (Figure 5b & 5d; p < 0.05). As seen in Figure 5b & 5d, GSR did not have any significant effect on the similarity between the EEG-related connectivity signatures and fMRI connectivity matrices, neither for alpha nor beta power variations.

## 4. Discussion

GSR is a widely used preprocessing step to remove global artifacts (mostly heart rate and breathing effects, collectively termed SLFOs) from fMRI data (Ciric et al., 2017; Parkes et al., 2018; Power et al., 2018, 2017b; Kassinopoulos & Mitsis, 2021), but remains controversial because it may also discard neural signals (Liu et al., 2017; Murphy and Fox, 2017). Up to date, the vast majority of studies investigating the origins of the global signal and evaluating the potential effects of GSR have examined fMRI data concurrently with either physiological or electrophysiological recordings, but not both at the same time. In the present study, we used simultaneous EEG-fMRI data, as well as cardiac and breathing recordings, to examine the processes underpinning the global signal and the impact of GSR on measures of brain activity and connectivity related to neural and systemic physiological fluctuations. Our results suggest that the global signal is strongly associated with physiological processes (R ∼ 0.6) and more weakly associated with EEG power fluctuations (R ∼ -0.1). We further demonstrated that GSR effectively removes the connectome patterns induced by physiological processes (SLFOs), but preserves the connectome patterns associated with EEG alpha and beta power fluctuations. These results provide evidence that, in the context of connectivity analyses, GSR improves the denoising of fMRI data and does not seem to alter the connectivity profiles associated with electrophysiological activity.

### 4.1 Association of SLFOs and EEG power with the fMRI global signal

We first evaluated the unique contributions of SLFOs and EEG activity to the global signal. SLFOs were found to explain a considerable fraction of global signal variance at rest (Figure 3a,3c), consistent with several earlier studies (Birn et al., 2006; Chang and Glover, 2009; Erdoğan et al., 2016; Kassinopoulos and Mitsis, 2019; Power et al., 2017b), as well as during the motor task (Figure 3b). Alpha power fluctuations were found to be negatively correlated with the global signal for all scans (Figure 3a-c), consistent with earlier work (Wong et al., 2013; Liu et al., 2017; Han et al., 2019), and beta power was found to be negatively correlated with the global signal during the motor task (Figure 3b). Our results suggest that contributions of electrophysiological origin to the global signal, albeit significant, were substantially weaker than physiological contributions.

A possible explanation for this is that the sign of correlation between fMRI fluctuations and EEG activity may not be the same across brain regions and, thus, when averaging fMRI signals across the whole brain, the contributions of these electrophysiological fluctuations may cancel out to some extent. Several studies have reported that, during resting conditions, alpha activity is negatively correlated with sensorimotor areas but positively correlated with default-mode regions (Mantini et al., 2007; Mayhew and Bagshaw, 2017; Scheeringa et al., 2012). These trends were also observed in our data, resulting in the presence of both positively and negatively correlated activity in Figure a. Therefore, it is indeed likely that the weak relationship between global signal and EEG activity may be due to differences in the polarity of associated fMRI activity across regions.

### 4.2 Association between SLFOs and alpha power

We observed that alpha power and SLFOs were negatively correlated during resting-state 1, albeit weakly. This finding was not replicated during the motor task and resting-state 2. The weak association between SLFOs and alpha activity may be due to the fact that subjects had their eyes open, which is in agreement to previous work that has demonstrated an association between respiration and alpha power during an eyes closed condition but not during eyes open (Yuan et al., 2013). Independent of this, shared contributions from SLFOs and alpha activity to the global signal, which could potentially reflect a direct or indirect effect of autonomic activity, were not consistently observed in our data. Therefore, our results seem to indicate that the major contributor to the global signal was physiological in origin.

### 4.3 Effect of GSR on connectome patterns associated with SLFOs and EEG power

We initially assessed the systematic effect of SLFOs on estimates of functional connectivity. The grand-averaged functional connectivity matrices calculated from the SLFOs synthetic datasets exhibited a heterogeneous pattern, characterized by stronger correlations within and between sensory cortices, including the visual cortex, somatosensory cortex and auditory cortex, as well as subcortical regions such as the thalamus and cerebellum (Figure 4a). This heterogeneity among brain regions is not surprising as it has been reported that global signal fluctuations are non-uniformly distributed across the brain (Fox et al., 2009; Kassinopoulos and Mitsis, 2019; Power et al., 2017b). Furthermore, the artifactual connectivity patterns due to SLFOs observed in the present study were similar to those reported in our recent study where a large number of healthy subjects from the Human Connectome Project (HCP) dataset was used for cortical regions (Xifra-Porxas et al., 2020). However, our results also revealed artifactual connectivity patterns within and between subcortical regions such as the thalamus and cerebellum, which we were unable to observe in the HCP dataset, likely due to poor signal-to-noise ratio in the subcortex in the HCP data (Ji et al., 2018; Seitzman et al., 2020).

Importantly, we observed a task- and session-related modulation of SLFOs-induced connectivity patterns (Figure 4a). Specifically, contributions of SLFOs were weaker during the motor task and the second resting-state period compared to the first. This observation may seem paradoxical as the contribution of SLFOs to the global signal did not significantly decrease across scans (Figure 3), but could be explained by the fact that global signal fluctuations were found to be reduced during the motor task and the second resting-state period compared to the first resting-state period (Supp. Fig. 3). It may also indicate that subjects were more alert during and after the motor task (Wong et al., 2013; Yeo et al., 2015). This interpretation is further supported by the decreased variability in heart rate observed during the second resting-state period compared to the first (p = 0.04; Supp. Fig. 5), suggesting a possible reduction in autonomic fluctuations. Finally, it could reflect the effect of circadian/ultradian physiological fluctuations, given that scanning started at 6 pm for all subjects and it has been reported that the amplitude of global signal fluctuations decreases as the day progresses (Orban et al., 2020).

The grand-averaged functional connectivity matrices calculated from the fraction of BOLD variance explained by alpha and beta power fluctuations also revealed structured patterns (Figure 5a, 5c). At rest, fluctuations in alpha power were associated with correlated activity within and between visual and dorsal attention networks (Figure 5a). Fluctuations in alpha power were also associated with anticorrelations of visual and dorsal attention networks with the default mode network and several subcortical regions. These results are consistent with earlier work reporting a positive association of alpha power fluctuations with fMRI activity in the default mode network and several subcortical regions, as well as a negative association between alpha power and fMRI activity in sensory regions (Bowman et al., 2017; Jann et al., 2009; Mantini et al., 2007; Mayhew and Bagshaw, 2017; Mo et al., 2013; Moosmann et al., 2003; Scheeringa et al., 2012). During the motor task, we found that alpha power was positively correlated with fMRI connectivity within the default mode network (Figure 5a), possibly reflecting the analogy between task-related synchronization/desynchronization alpha patterns and activation/deactivation default mode fluctuations (Mayhew et al., 2013; Mo et al., 2013). Furthermore, fluctuations in beta power were mostly associated with positively and negatively correlated activity within the visual and default mode networks, but surprisingly not in the somatosensory network (Figure 5**c**).

Regarding the effect of GSR on the estimates of functional connectivity, our recent work (Xifra-Porxas et al., 2020) provided evidence that GSR is effective in terms of removing systematic biases on measures of resting-state functional connectivity arising due to SLFOs. However, since we did not have electrophysiological recordings in that study, we were unable to assess whether GSR also removed any signal of interest. Here, we show that, in addition to mitigating the effect of sLFOs on resting-state connectivity patterns, GSR yielded similar results during the motor task (Figure 4b). It is worth noting that even though we used a relatively aggressive preprocessing pipeline (10 PCA components from white matter) (Kassinopoulos and Mitsis, 2022), performing GSR was found to be beneficial. Considering that in a large number of fMRI studies only the average signals from white matter and cerebrospinal fluid compartments are regressed out, our results suggest that the effectiveness of GSR with respect to artifact removal would have been even more pronounced for milder preprocessing pipelines. Furthermore, we found that EEG power fluctuations within the alpha and beta bands were correlated to the global signal (Figure 3) but the extent of shared variance was small and, thus, GSR did not have any significant effect on the fMRI connectivity patterns attributed to alpha (Figure 5b) and beta (Figure 5d) electrophysiological power, neither at rest nor during the motor task. This is to some extent supported by the findings reported in a study involving macaque monkeys, where it was shown that, while inactivation of the basal forebrain led to selective suppression of ipsilateral global components, resting-state brain networks preserved their distinctive topography (Turchi et al., 2018). Overall, our results support the view that the benefits of GSR in terms of fMRI denoising outweigh the potential loss of neural information (Li et al., 2019a, 2019b).

### 4.4 Limitations

A key limitation of the present study is the relatively small sample size (n = 11), which we acknowledge may constrain the generalizability of our findings and may also contribute to the non-significant effects of GSR on the connectivity patterns related to neural activity. However, the primary aim of this work was to establish a methodological proof-of-concept regarding the differential impact of global signal regression on connectivity patterns linked to physiological signals and neural oscillations. Despite the known inter-subject variability in EEG-fMRI studies, the consistency of our results across subjects and scanning sessions suggests that the observed effects are robust. Notably, we were able to replicate previous findings from a large cohort with regards to the effect of GSR on connectivity patterns induced by SLFOs (Xifra-Porxas et al., 2021), which lends support to the validity of our results. Further research with larger sample sizes will be essential to provide further insights into the nuanced impact of GSR on neural connectivity patterns.

Furthermore, despite the large fraction of the global signal variance that was explained by SLFOs, and to a lesser extent by EEG power (Figure 3), there was still variance unaccounted for. The EEG signal is not a perfect reflection of neural activity and is known to be considerably noisy in the MR environment, as well as mostly blind to deeper sources. Therefore, it is likely that there was an additional fraction of neural activity that was present in the global signal that we were unable to detect in the EEG data. Likewise, there are other physiological factors not considered here that are known to give rise to SLFOs and are reflected on the global signal, such as finger skin vascular tone (Kassinopoulos and Mitsis, 2021; Özbay et al., 2019), arterial CO2 changes (Prokopiou et al., 2019; Wise et al., 2004) and arterial blood pressure (Dagenais and Mitsis, 2023; Whittaker et al., 2019b). Future studies using more direct surrogates of neuronal activity (e.g. intracranial EEG) are needed to confirm whether crucial neural information is being removed through GSR.

Another limitation concerns the simplicity of the EEG indices used in this study. We focused on global EEG power averaged across channels to remain consistent with prior work (e.g., Wong et al., 2013; 2016), and to avoid increasing model complexity given our modest sample size. However, we acknowledge that more spatially resolved EEG metrics, such as alpha asymmetry or region-specific alpha spindles, have been widely applied in cognitive and affective neuroscience (Ismail and Karwowski, 2020; Giannakakis et al., 2022), and may reveal stronger associations with the fMRI global signal. These metrics have not yet been systematically examined in resting-state EEG-fMRI studies. Future research with larger cohorts and improved spatial modeling will be needed to clarify their relevance to global signal fluctuations.

Moreover, while our results suggest that GSR effectively mitigates physiological confounds without substantially altering EEG-derived connectivity patterns, we cannot exclude the possibility that GSR also suppresses neural signal components that are spatially widespread or temporally synchronized across regions. Prior studies have shown that GSR can introduce anti-correlations and reduce the amplitude of functional connectivity estimates (Murphy et al., 2009), and that the global signal may contain meaningful neural fluctuations related to arousal and vigilance (Liu et al., 2017). Therefore, it remains possible that subtle neural contributions to the BOLD signal, particularly those not captured by global EEG power, are attenuated. Future studies incorporating more sensitive electrophysiological markers, such as phase-based metrics or cross-frequency coupling, may help clarify the extent to which GSR affects neural signal components.

Finally, we examined resting conditions and a hand-grip task, but results could be different during other conditions, such as sleep (Duyn et al., 2020) or when investigating diurnal variations in large-scale spontaneous brain activity (Orban et al., 2020).

## 5. Conclusions

Using simultaneous EEG-fMRI recordings, we demonstrated that the global BOLD fMRI signal exhibits a significant association with SLFOs as well as EEG alpha and beta power fluctuations, during resting conditions and motor task execution. In both cases, the effect of SLFOs was found to be more pronounced. We also provide evidence that GSR effectively removes confounds in functional connectivity induced by SLFOs, without severely disrupting the functional connectivity patterns attributed to EEG alpha and beta activity, suggesting that it overall yields a beneficial effect in terms of fMRI-based connectivity assessment.

## Funding

This work was supported by funds from Fonds de la Recherche du Québec – Nature et Technologies (FRQNT) Team Grants 2016-191780 [MHB & GDM] and 2018-256480 [GDM], the Natural Sciences and Engineering Research Council of Canada (NSERC) Discovery Grant RGPIN-2019-06638 [GDM], the Canadian Foundation for Innovation grant numbers 34277 [MHB] and 34362 [GDM], and the Canada First Research Excellence Fund (awarded to McGill University for the Healthy Brains for Healthy Lives (HBHL) initiative) [MK]. AXP and MK received financial support through funding from McGill University and the Québec Bio-imaging Network (QBIN).

## Acknowledgments

We would like to thank the MRI technicians at the Montreal Neurological Institute for their patience and assistance collecting the data.

## Declaration of conflicting interests

The authors declared no potential conflicts of interest with respect to the research, authorship, and/or publication of this article.

## Statements and Declarations

Not applicable.

## Authors’ contributions

AXP: conceptualization, study design, participant recruitment, data acquisition, preprocessing, formal analysis, figure design, writing & revision; MK: conceptualization, experimental paradigm implementation, data acquisition setup, data acquisition, preprocessing assistance, result interpretation, revision; PP: data acquisition, preprocessing assistance, result interpretation; MHB: study design, revision. GDM: conceptualization, supervision, revision.

## Data availability statement

The data that support the findings of this study are available from the corresponding author upon reasonable request.

## Ethical Approval and Informed Consent

The study was approved by the McGill University Ethical Advisory Committee. All participants provided written informed consent before participation and were compensated for their participation.

## Supplementary Material

Supplementary material for this article has been submitted and will be available online upon publication.

## Notes

### Competing Interest Statement

The authors have declared no competing interest.

### Summary of Updates

The manuscript title was revised to a more declarative form: Global signal regression reduces connectivity patterns related to physiological signals and does not alter EEG-derived connectivity.

