## Supplementary Material for "Global signal regression reduces connectivity patterns related to physiological signals and does not alter EEG-derived connectivity"

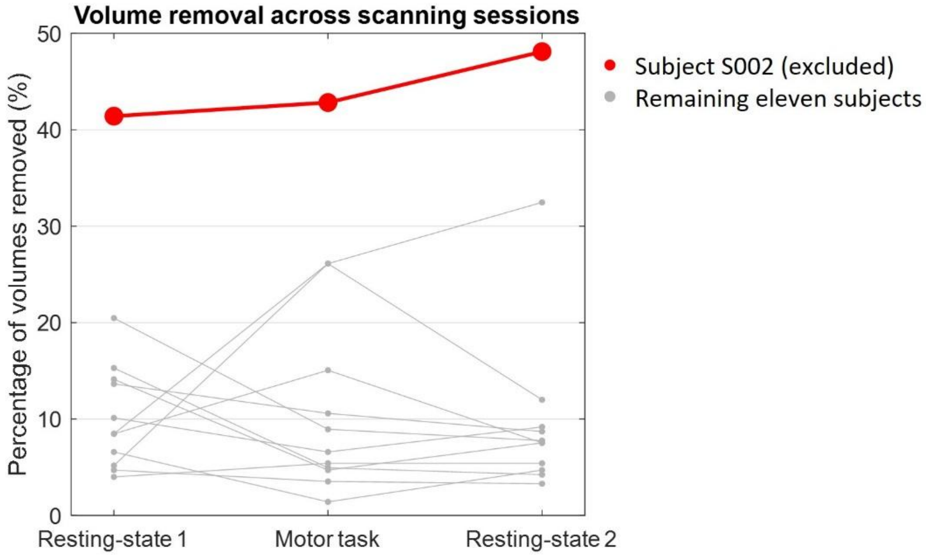


**Supp. Fig. 1.** Percentage of volumes removed per subject across the three scanning sessions. Subject S002 was excluded from further analysis due to consistently high motion-related volume removal (>40%) in all three scanning sessions, resulting in a final sample of eleven subjects.

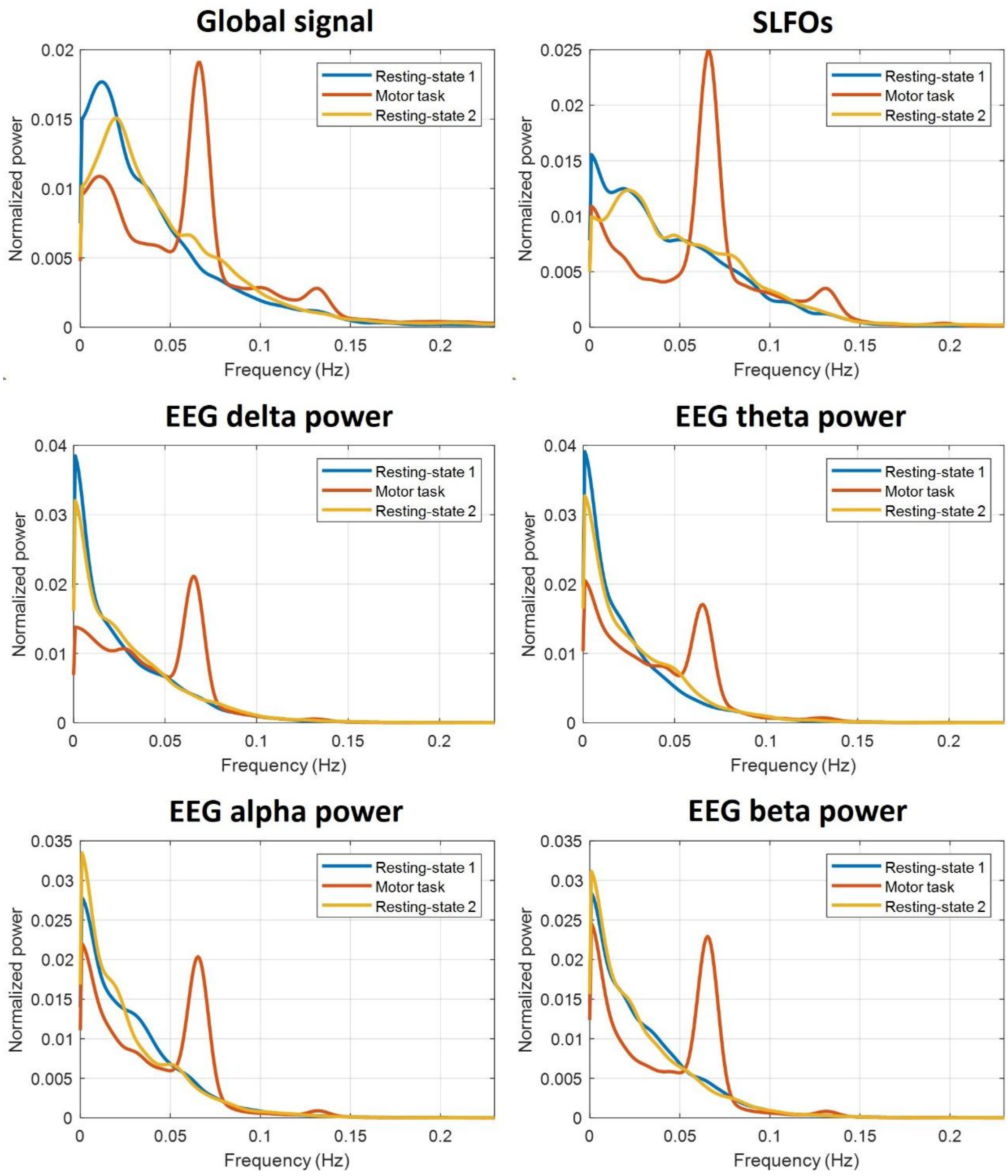


**Supp. Fig. 2.** Power spectral density of the fMRI global signal, the SLFOs, and the EEG-derived band power signals (delta, theta, alpha, beta), averaged across all subjects. Spectra were computed using Welch's method and normalized to unit area for comparability. The global signal exhibited dominant power in the low-frequency range (<0.15 Hz) across all three scanning sessions, with a distinct peak near 0.07 Hz during the motor task, which was associated with the task paradigm. SLFOs showed consistent spectral profiles across sessions. The EEG band power signals displayed similar spectral shapes, with their energy concentrated in slightly lower frequencies.

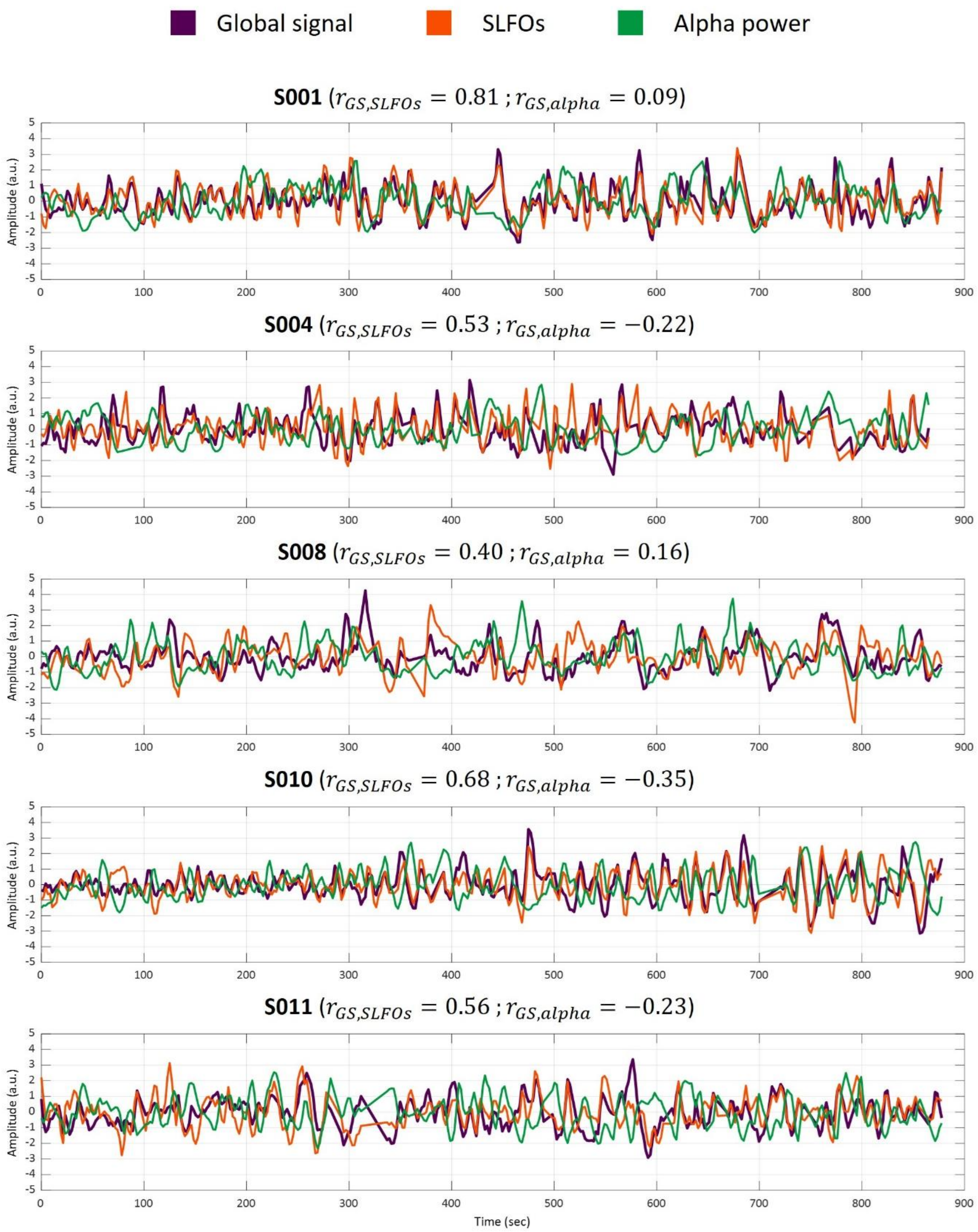


**Supp. Fig. 3.** Global signal, SLFOs, and alpha power traces from 5 representative subjects.

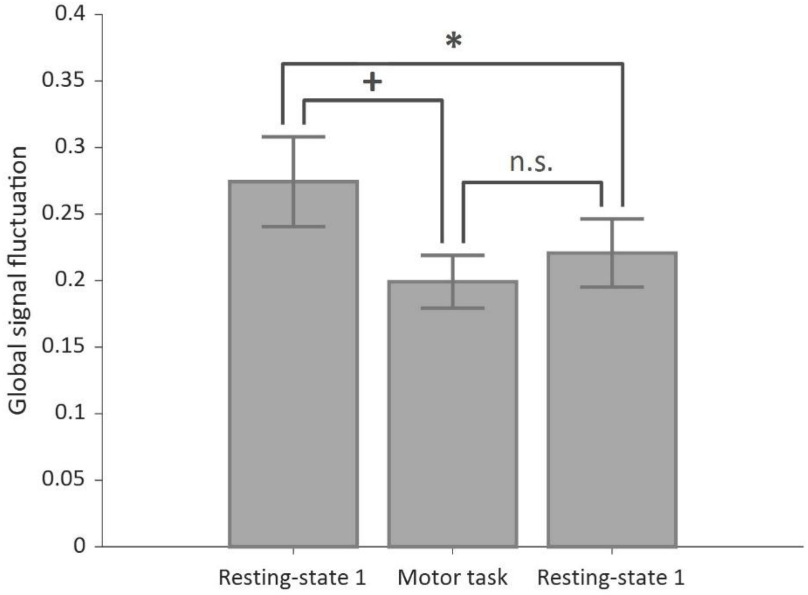


**Supp. Fig. 4.** Mean global signal fluctuation (standard deviation of the global signal) across participants for each scan. Error bars denote standard error of the mean. Significance testing using the sign-rank test (* $p$ <0.05, ⁺ $p$ <0.1).

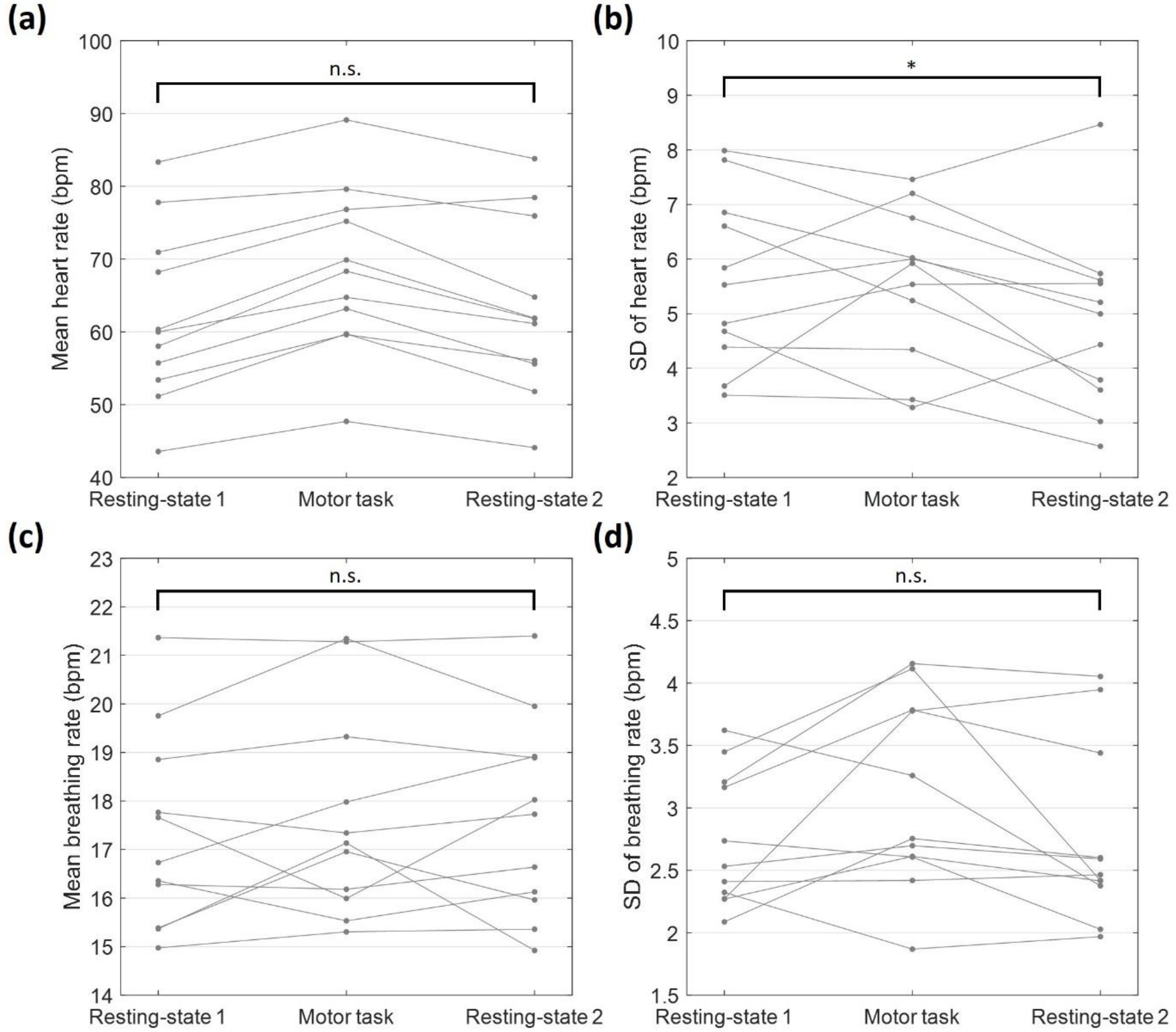


**Supp. Fig. 5.** Heart rate and breathing rate metrics across participants for each scanning session. (a) Mean heart rate, (b) standard deviation (SD) of heart rate, (c) mean breathing rate, and (d) SD of breathing rate across the three scanning sessions. Significance testing was performed using the sign-rank test (* p <0.05).
